# Protein-coding variation and introgression of regulatory alleles drive plumage pattern diversity in the rock pigeon

**DOI:** 10.1101/242552

**Authors:** Anna I. Vickrey, Rebecca Bruders, Zev Kronenberg, Emma Mackey, Ryan J. Bohlender, Emily T. Maclary, E.J. Osborne, Kevin P. Johnson, Chad D. Huff, Mark Yandell, Michael D. Shapiro

## Abstract

Birds and other vertebrates display stunning variation in pigmentation patterning, yet the genes controlling this diversity remain largely unknown. Rock pigeons (*Columba livia*) are fundamentally one of four color pattern phenotypes, in decreasing order of melanism: T-check, checker, bar (ancestral), or barless. Using whole-genome scans, we identified *NDP* as a candidate gene for this variation. Allele-specific expression differences in *NDP* indicate *cis*-regulatory differences between ancestral and melanistic alleles. Sequence comparisons suggest that derived alleles originated in the speckled pigeon (*Columba guinea*), providing a striking example of introgression of alleles that are favored by breeders and are potentially advantageous in the wild. In contrast, barless rock pigeons have an increased incidence of vision defects and, like two human families with hereditary blindness, carry start-codon mutations in *NDP*. In summary, we find unexpected links between color pattern, introgression, and vision defects associated with regulatory and coding variation at a single locus.

## INTRODUCTION

Vertebrates have evolved a vast array of epidermal colors and color patterns in response to natural, sexual, and artificial selection. Numerous studies have identified key genes that determine variation in the types of pigments that are produced (e.g., Hubbard et al. 2010; Manceau et al. 2010; Roulin and Ducrest 2013; Domyan et al. 2014; Rosenblum et al. 2014). In contrast, considerably less is known about the genetic mechanisms that determine pigment *patterning* throughout the entire epidermis and within individual epidermal appendages (e.g., feathers, scales, and hairs) (Kelsh 2004; Protas and Patel 2008; Kelsh et al. 2009; Lin et al. 2009; Kaelin et al. 2012; Lin et al. 2013; Eom et al. 2015; Poelstra et al. 2015; Mallarino et al. 2016). In birds, color patterns are strikingly diverse among different populations and species, and these traits have profound impacts on mate-choice, crypsis, and communication (Hill and McGraw 2006).

The domestic rock pigeon (*Columba livia*) displays enormous phenotypic diversity within and among over 350 breeds, including a wide variety of plumage pigmentation patterns (Shapiro and Domyan 2013; Domyan and Shapiro 2017). Some of these pattern phenotypes are found in feral and wild populations as well (Johnston and Janiga 1995). A large number of genetic loci contribute to pattern variation in rock pigeons, including genes that contribute in an additive fashion and others that epistatically mask the effects of other loci (Jones 1922; Hollander 1937; Sell 2012; Domyan et al. 2014). Despite the genetic complexity of the full spectrum of plumage pattern diversity in pigeons, classical genetic experiments demonstrate that major wing shield pigmentation phenotypes are determined by an allelic series at a single locus (*C*, for “checker” pattern) that produces four phenotypes: T-check (*C^T^* allele, also called T-pattern), checker (C), bar (+), and barless (*c*), in decreasing order of dominance and melanism (Fig. 1A) (Bonhote and Smalley 1911; Hollander 1938; Hollander 1983; Levi 1986; Sell 2012). Bar is the ancestral phenotype (Darwin 1859, 1868), yet checker and T-check can occur at higher frequencies than bar in urban feral populations, suggesting a fitness advantage in areas of dense human habitation (Obukhova and Kreslavskii 1984; Johnston and Janiga 1995; Čanády and Mošanský 2013).

**Fig. 1.**
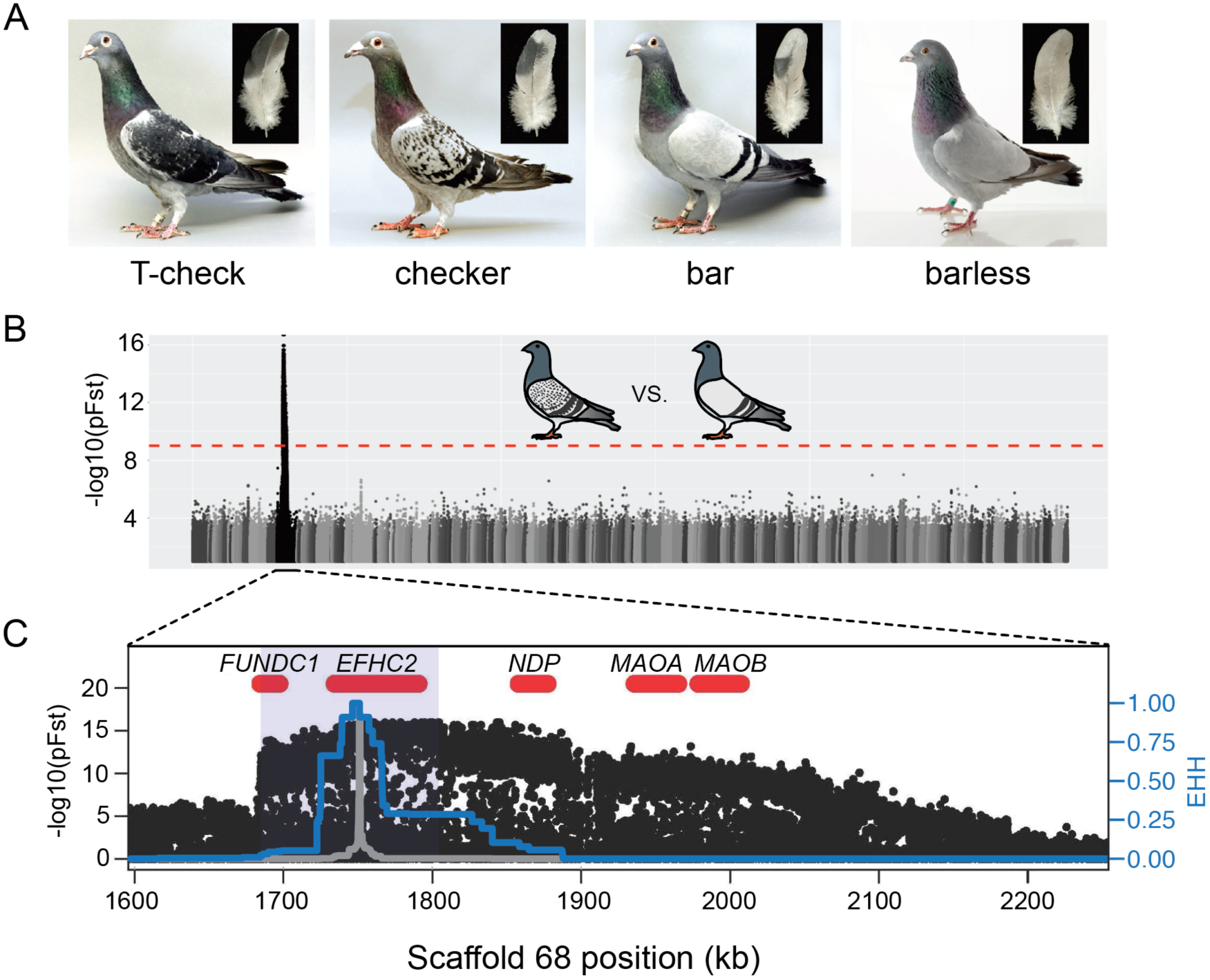
A single genomic region is associated with rock pigeon (*C. livia*) wing pigmentation pattern (A) Four classical wing pattern pigmentation phenotypes, shown in decreasing order of genetic dominance and melanism (left to right): T-check, checker, bar, and barless. Photos courtesy of the Genetics Science Learning Center (http://learn.genetics.utah.edu/content/pigeons). (B) Whole-genome pFst comparisons between the genomes of bar (n=17) and checker (n=24) pigeons. Dashed red line marks the genome-wide significance threshold (9.72e-10). (C) Detail of pFst peak shows region of high differentiation on Scaffold 68. Five genes within the region are shown in red. Blue shading marks the location of the smallest shared haplotype common to all checker and T-check birds. Haplotype homozygosity in the candidate region extends further for checker and t-check birds (blue trace) than for bar birds (gray), a signature of positive selection for the derived alleles. Extended haplotype homozygosity (EHH) was measured from focal position 1,751,072 and follows the method of Sabeti et al. (2007).

Color pattern variation is associated with several important life history traits in feral pigeon populations. For example, checker and T-check birds have higher frequencies of successful fledging from the nest, longer (up to year-round) breeding seasons, and can sequester more toxic heavy metals in plumage pigments through chelation (Petersen and Williamson 1949b; Lofts et al. 1966; Murton et al. 1973; Chatelain et al. 2014; Chatelain et al. 2016). Relative to bar, checker and T-check birds also have reduced fat storage and, perhaps as a consequence, lower overwinter adult survival rates in harsh rural environments (Petersen and Williamson 1949a; Jacquin et al. 2012). Disassortative mating occurs in feral pigeons with different patterns, so sexual selection probably influences the frequencies of wing pigmentation patterns in feral populations as well (Burley 1981; Johnston and Johnson 1989). In contrast, barless, the recessive and least melanistic phenotype, is rarely observed in feral pigeons (Johnston and Janiga 1995). In domestic populations, barless birds have a higher frequency of vision defects, sometimes referred to as “foggy” vision (Hollander and Miller 1981; Hollander 1983; Mangile 1987), which could negatively impact fitness in the wild.

In this study, we investigate the molecular and evolutionary mechanisms underlying wing pattern diversity in pigeons. We discover both coding and regulatory variation at a single candidate gene, and a trans-species polymorphism linked with pattern variation within and between species that likely resulted from interspecies hybridization.

## RESULTS AND DISCUSSION

### A genomic region on Scaffold 68 is associated with wing pattern phenotype

To identify the genomic region containing the major wing pigmentation pattern locus, we used a probabilistic measure of allele frequency differentiation (pFst; Domyan et al. 2016) to compare the resequenced genomes of bar pigeons to genomes of pigeons with either checker or T-check patterns (see Methods). Checker and T-check birds were grouped together because these two patterns are sometimes difficult to distinguish, even for experienced hobbyists (Fig. 1A). Checker birds are typically less pigmented than T-check birds, but genetic modifiers of pattern phenotypes can minimize this difference. A two-step whole-genome scan (see Methods, Fig. 1B, Fig. S1A) identified a single ~103-kb significantly differentiated region on Scaffold 68 that was shared by all checker and T-check birds (position 1,702,691–1,805,600 of the Cliv_1.0 pigeon genome assembly (Shapiro et al. 2013); p = 1.11e-16, genome-wide significance threshold = 9.72e-10). The minimal shared region was defined by haplotype breakpoints in a homozygous checker and a homozygous bar bird and is hereafter referred to as the minimal checker haplotype. As expected for the well-characterized allelic series at the *C* locus, we also found that a broadly overlapping region of Scaffold 68 was highly differentiated between the genomes of bar and barless birds (p = 3.11e-15, genome-wide significance threshold = 9.71e-10; Fig. S1B). Together, whole-genome comparisons identified a single genomic region corresponding to the wing pattern *C* locus.

### A copy number variant is associated with melanistic wing patterns

To identify genetic variants associated with the derived checker and T-check phenotypes, we first compared annotated protein-coding genes throughout the genome. We found a single, predicted, fixed change in EFHC2 (Y572C, Fig. S2) in checker and T-check birds relative to bar birds (VAAST; Yandell et al. 2011). However, this same amino acid substitution is also found in *Columba rupestris*, a closely related species to *C. livia* that has a bar wing pattern. Thus, the Y572C substitution is not likely to be causative for the checker or T-check pattern, nor is it likely to have a strong impact on protein function (MutPred2 score 0.468, no recognized affected domain; PolyPhen-2 score 0.036; Adzhubei et al. 2010; Pejaver et al. 2017).

Next, we examined sequence coverage across the checker haplotype and discovered a copy number variable (CNV) region (approximate breakpoints at Scaffold 68 positions 1,790,000 and 1,805,600). Based on normalized read-depths of resequenced birds, we determined that the CNV region has one, two, or four copies per chromosome. Bar birds (n=12) in our resequencing panel always had a total of two copies in the CNV region (one on each chromosome), but most checker (n=5 of 7) and T-check (n=2 of 2) genomes examined had additional copies of the CNV (Fig. 2A). Using a PCR assay to amplify across the breakpoints in birds with more than one copy per chromosome, we determined that additional copies result from tandem repeats. We found no evidence that the checker haplotype contains an inversion based on mapping of paired-end reads at the CNV breakpoints (WHAM; Kronenberg et al. 2015). In addition, we were able to amplify unique PCR products that span the outer CNV breakpoints (data not shown), suggesting that there are no inversions within the CNV region.

**Fig. 2.**
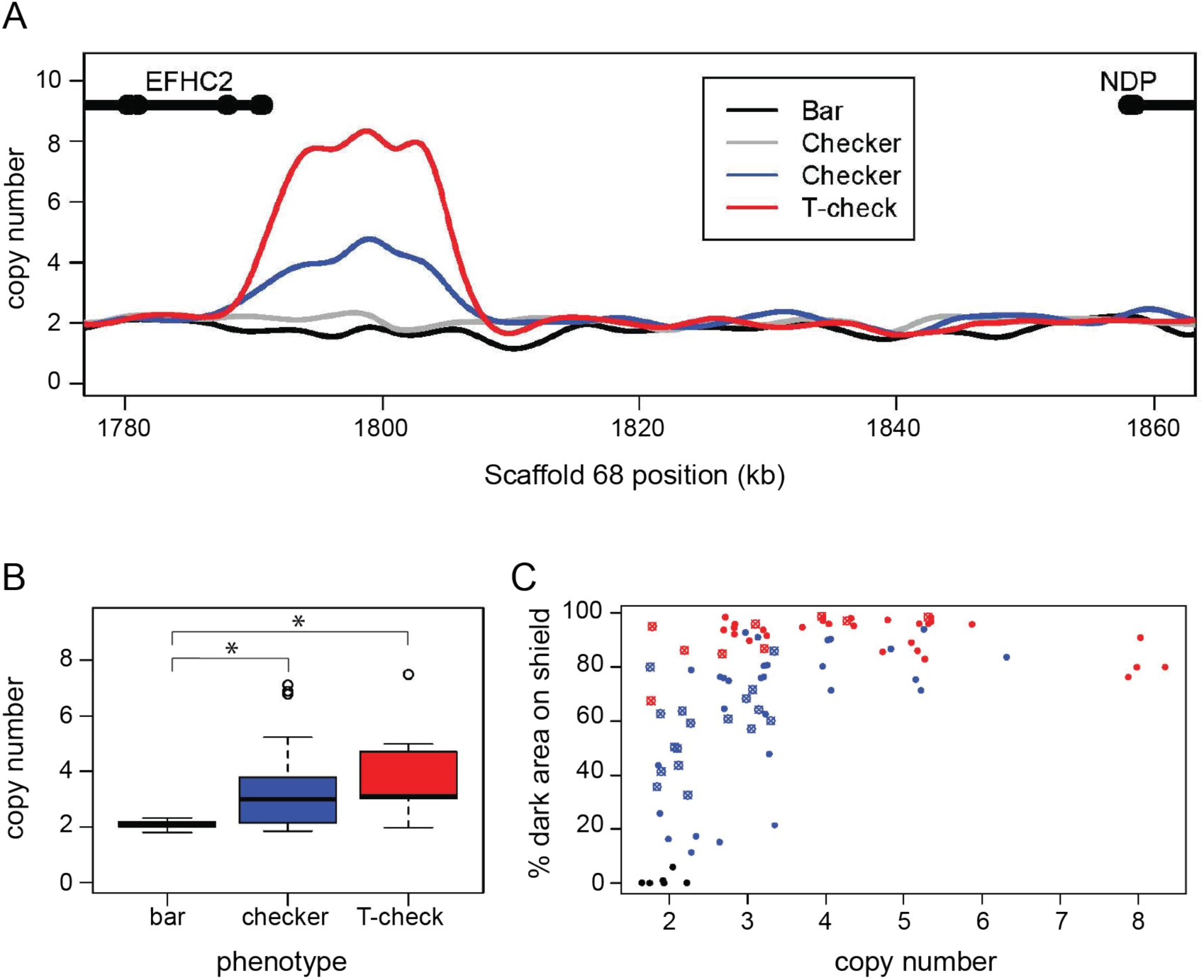
A copy number variant (CNV) in the candidate region is associated with T-check and checker phenotypes. (A) Normalized read depths from resequenced birds are plotted in the candidate region between *EFHC2* and *NDP* on Scaffold 68. Thickened portions of gene models represent exons and thin portions are introns. Representative individual read depth traces are shown for the following: black for bar *C. livia*, grey for checker *C. livia* individuals without additional copies of the CNV, blue for checker *C. livia* individuals with additional copies of the CNV region, red for T-check *C. livia*. (B) CNV quantification for 94 birds (20 bar, 56 checker, and 18 T-check). Checker and T-check phenotypes (as reported by breeders) were associated with increased copy numbers (p=2.1e-05). (C) CNV and phenotype quantification for an additional 84 birds, including 26 feral pigeons. Increased copy number was associated with an increase in dark area on the wing shield (r^2^=0.46, linear regression). Points are colored by reported phenotype and origin: bar, black; checker, blue; T-check, red; domestic breeds, solid points; ferals, cross points.

The fact that some checker birds had only two total copies of the CNV region demonstrates that a copy number increase is not necessary to produce melanistic phenotypes. However, consistent with the dominant inheritance pattern of the phenotype, all checker and T-check birds had at least one copy of the checker haplotype. Thus, a checker haplotype on at least one chromosome appears to be necessary for the dominant melanistic phenotypes, but additional copies of the CNV region are not.

In a larger sample of pigeons, we found a significant association between copy number and phenotype (TaqMan assay; pairwise Wilcoxon test, p=2.1e-05). Checker (n=40 of 56) and T-check (n=15 of 18) patterns were associated with additional copies, and pigeons with the bar pattern (n=20) never had more than two copies in total (Fig. 2B). Although additional copies of the CNV only occurred in checker and T-check birds, we did not observe a consistent number of copies associated with either phenotype. This could be due to a variety of factors, including modifiers that darken genotypically-checker birds to closely resemble T-check (Jones 1922; Sell 2012) and environmental factors such as temperature-dependent darkening of the wing shield during feather development (Podhradsky 1968).

Due to the potential ambiguity in categorical phenotyping, we next measured the percent of pigmented area on the wing shield and tested for associations between copy number and percentage of pigmented wing shield area. We phenotyped and genotyped an additional 63 birds from diverse domestic breeds as well as 26 feral birds, and found that estimated copy number in the variable region was correlated with the amount of dark pigment on the wing shield (nonlinear least squares regression, followed by r^2^ calculation; r^2^=0.46) (Fig. 2C). This correlation was a better fit to the regression when ferals were excluded (r^2^=0.68, Fig. S3B), possibly because numerous pigmentation modifiers (e.g., *sooty* and *dirty*) are segregating in feral populations (Hollander 1938; Johnston and Janiga 1995). Together, our analyses of genetic variation among phenotypes point to a CNV that is associated with qualitative and quantitative color pattern variation in pigeons.

### *NDP* is differentially expressed in feather buds of different wing pattern phenotypes

The CNV that is associated with wing pattern variation resides between two genes, *EFHC2* and *NDP. EFHC2* is a component of motile cilia, and mouse mutants have juvenile myoclonic epilepsy (Linck et al. 2014). In humans, allelic variation in *EFHC2* is also associated with differential fear responses and social cognition (Weiss et al. 2007; Blaya et al. 2009; Startin et al. 2015; but see Zinn et al. 2008). However, *EFHC2* has not been implicated in pigmentation phenotypes in any organism. *NDP* encodes a secreted ligand that activates *WNT* signaling by binding to its only known receptor FZD4 and its co-receptor LRP5 (Smallwood et al. 2007; Hendrickx and Leyns 2008; Deng et al. 2013; Ke et al. 2013). Notably, *NDP* is one of many differentially expressed genes in the feathers of closely related crow subspecies that differ, in part, by the intensity of plumage pigmentation (Poelstra et al. 2015). Furthermore, FZD4 is a known melanocyte stem cell marker (Yamada et al. 2010). Thus, based on expression variation in different crow plumage phenotypes, and the expression of its receptor in pigment cell precursors, *NDP* is a strong candidate for pigment variation in pigeons.

The CNV in the intergenic space between *EFHC2* and *NDP* in the candidate region, coupled with the lack of candidate coding variants between bar and checker haplotypes, led us to hypothesize that the CNV region might contain regulatory variation that could alter expression of one or both neighboring genes. To test this possibility, we performed qRT-PCR on RNA harvested from regenerating wing shield feathers of bar, checker, and T-check birds. *EFHC2* was not differentially expressed between bar and either checker or T-check patterned feathers (p=0.19, pairwise Wilcoxon test, p-value adjustment method: fdr), although expression levels differed slightly between the checker and T-check patterned feathers (p=0.046, Fig. 3A). Expression levels of other genes adjacent to the minimal checker haplotype region also did not vary by phenotype (Fig. S4).

**Fig. 3.**
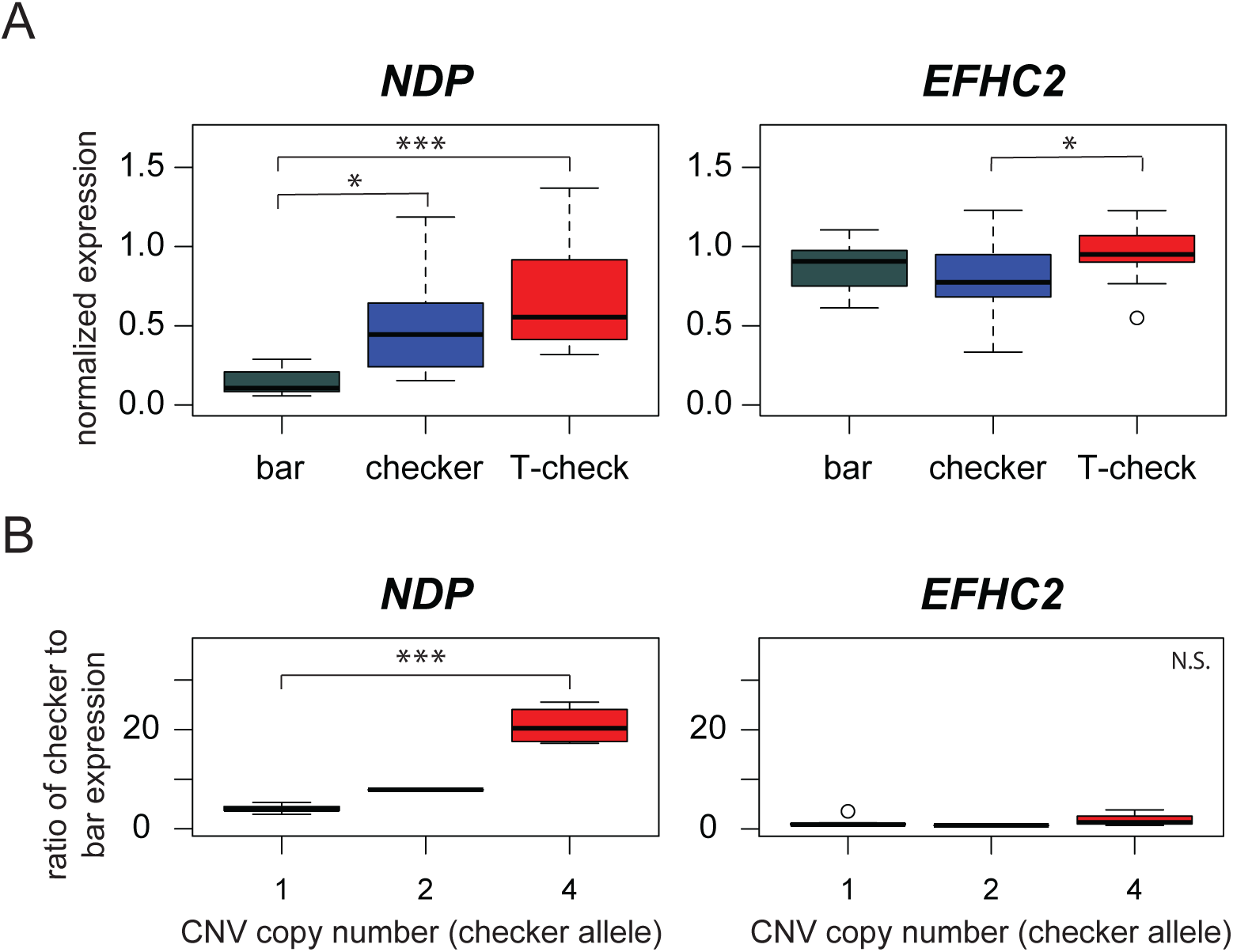
Expression differences in *NDP*, but not *EFHC2*, indicate *cis*-regulatory differences associated with pigmentation phenotypes. (A) qRT-PCR assays demonstrate higher expression of *NDP* in regenerating feathers of checker and T-check birds than in bar birds. Expression levels of *EFHC2* are indistinguishable between bar and melanistic phenotypes (p=0.19), although checker and T-check differed from each other (p=0.046). (B) Allele-specific expression assay in regenerating feathers from heterozygous bar/checker birds for *NDP* and *EFHC2*. Copies of the CNV region on the checker chromosome were quantified using a custom Taqman assay. Boxes span the first to third quartiles, bars extend to minimum and maximum observed values, black line indicates median. Expression of *EFHC2* alleles were not significantly different, and checker alleles of *NDP* showed higher expression than the bar allele; p=0.0028 for two-sample t-test between 1 vs. 4 copies, p=1.84e-06 for glm regression.

In contrast, expression of *NDP* was significantly increased in checker feathers – and even higher in T-check feathers – relative to bar feathers (Fig. 3A) (bar-checker comparison, p=1.9e-05; bar-T-check, p=1.0e-08; checker-T-check, p=0.0071; pairwise Wilcoxon test, all comparisons were significant at a false discovery rate of 0.05). Moreover, when qRT-PCR expression data for checker and T-check feathers were grouped by copy number instead of categorical phenotype, the number of CNV copies was positively associated with *NDP* expression level (Fig. S5). Thus, expression of *NDP* is positively associated with both increased melanism (categorical pigment pattern phenotype) and CNV genotype.

The increase in *NDP* expression could be the outcome of at least two molecular mechanisms. First, one or more regulatory elements in the CNV region (or elsewhere) could increase expression of *NDP* in *cis*. Such changes would only affect expression of the allele on the same chromosome (Wittkopp et al. 2004). Second, *trans-acting* factors encoded within the minimal checker haplotype (e.g., *EFHC2* or an unannotated feature) could increase *NDP* expression, resulting in an upregulation of *NDP* alleles on both chromosomes.

To distinguish between these possibilities, we carried out allele-specific expression assays (Domyan et al. 2014; Domyan et al. 2016) on the regenerating feathers of birds that were heterozygous for bar and checker alleles in the candidate region (checker alleles with one, two, or four copies of the CNV). In the common trans-acting cellular environment of heterozygous birds, checker alleles of *NDP* were more highly expressed than bar alleles, and these differences were further amplified in checker alleles with more copies of the CNV (Fig. 3B) (p=0.0028 for two-sample t-test between 1 vs. 4 copies, p=1.84e-06 for generalized linear model regression). In comparison, transcripts of *EFHC2* from checker and bar alleles were not differentially expressed in the hybrid background (Fig. 3B) (p=0.5451 for two-sample t-test between 1 vs. 4 copies, p=0.471 for linear regression). Together, our expression studies indicate that a *cis*-acting regulatory change drives increased expression of *NDP* in pigeons with more melanistic plumage patterns, but does not alter expression of *EFHC2* or other nearby genes (Figs. 3A, S4). Furthermore, because *NDP* expression increases with additional copies of the CNV, the regulatory element probably resides within the CNV itself.

### A missense mutation at the start codon of *NDP* is associated with barless

In humans, mutations in *NDP* can result in Norrie disease, a recessively-inherited disorder characterized by a suite of symptoms including vision deficiencies, intellectual and motor impairments, and auditory deficiencies (Norrie 1927; Warburg 1961; Holmes 1971; Chen et al. 1992; Sims et al. 1992). Protein-coding mutations in *NDP*, including identical mutations segregating within single-family pedigrees, result in variable phenotypic outcomes, including incomplete penetrance (Meindl et al. 1995; Berger 1998; Allen et al. 2006). Intriguingly, barless pigeons also have an increased incidence of vision deficiencies and, as in humans with certain mutant alleles of *NDP*, this phenotype is not completely penetrant (Hollander 1983). Thus, based on the known allelism at the *C* locus, the nomination of regulatory changes at *NDP* as candidates for the *C* and *C^T^* alleles, and the vision-related symptoms of Norrie disease, *NDP* is also a strong candidate for the barless phenotype (*c* allele).

To test the prediction that variation in *NDP* is associated with the barless phenotype, we used VAAST to scan the resequenced genomes of 9 barless pigeons and found that all were homozygous for a nonsynonymous protein-coding change at the start codon of *NDP* that was perfectly associated with the barless wing pattern phenotype. We detected no other genes with fixed coding changes or regions of significant allele frequency differentiation (pFst) elsewhere in the genome. We genotyped an additional 14 barless birds and found that all were homozygous for the same start-codon mutation (Fig. S6). The barless mutation is predicted to truncate the amino terminus of the NDP protein by 11 amino acids, thereby disrupting the 24-amino acid signal peptide sequence (www.uniprot.org, Q00604 NDP_Human). *NDP* is still transcribed and detectable by RT-PCR in regenerating barless feathers (data not shown); therefore, we speculate that the start-codon mutation might alter the normal secretion of the protein into the extracellular matrix (Gierasch 1989).

In humans, coding mutations in *NDP* are frequently associated with a suite of neurological deficits. In pigeons, however, only wing pigment depletion and vision defects are reported in barless homozygotes. Remarkably, two human families segregating Norrie disease have only vision defects, and like barless pigeons, these individuals have start-codon mutations in *NDP* (Fig. S6) (Isashiki et al. 1995). Therefore, signal peptide mutations might affect a specific subset of developmental processes regulated by *NDP*, while leaving other (largely neurological) functions intact. In summary, wing pattern phenotypes in pigeons are associated with the evolution of both regulatory (checker, T-check) and coding (barless) changes in the same gene, and barless pigeons share a partially-penetrant visual deficiency with human patients who have start-codon substitutions.

### Signatures of introgression of the checker haplotype

Pigeon fanciers have long hypothesized that the checker pattern in the rock pigeon (*Columba livia*) resulted from a cross-species hybridization event with the speckled pigeon (*Columba guinea*, Fig. S7), a species with a checker-like wing pattern (G. Hochlan, G. Young, personal communication) (Hollander 1983). Although *C. livia* and *C. guinea* diverged an estimated 4-5 million years ago, inter-species crosses can produce fertile hybrids (Whitman 1919; Irwin et al. 1936; Taibel 1949; Miller 1953). Moreover, hybrid F_1_ and backcross progeny between *C. guinea* and bar *C. livia* have checkered wings, much like *C. livia* with the *C* allele (Whitman 1919; Taibel 1949). Taibel (1949) showed that, although hybrid F_1_ females were infertile, two more generations of backcrossing to *C. livia* could produce checker offspring of both sexes that were fully fertile. In short, Taibel introgressed the checker trait from *C. guinea* into *C. livia* in just three generations.

To evaluate the possibility of an ancient cross-species introgression event, we sequenced an individual *C. guinea* genome to ~30X coverage and mapped the reads to the *C. livia* reference assembly. We calculated four-taxon *D*-statistics (“ABBA-BABA” test; Durand et al. 2011) to test for deviations from expected sequence similarity between *C. guinea* and *C. livia*, using a wood pigeon (*C. palumbus*) genome as an outgroup. In this case, the null expectation is that the *C* candidate region will be more similar between conspecific bar and checker *C. livia* than either will be to the same region in *C. guinea*. That is, the phylogeny of the candidate region should be congruent with the species phylogeny. However, we found that the *D*-statistic approaches 1.0 in the candidate region (n=10 each for bar and checker *C. livia*), indicating that checker *C. livia* are more similar to *C. guinea* than to conspecific bar birds in this region (Fig. 4A). The mean genome-wide *D*-statistic was close to zero (0.021), indicating that bar and checker sequences are more similar to each other throughout the genome than either one is to *C. guinea*.

**Fig. 4.**
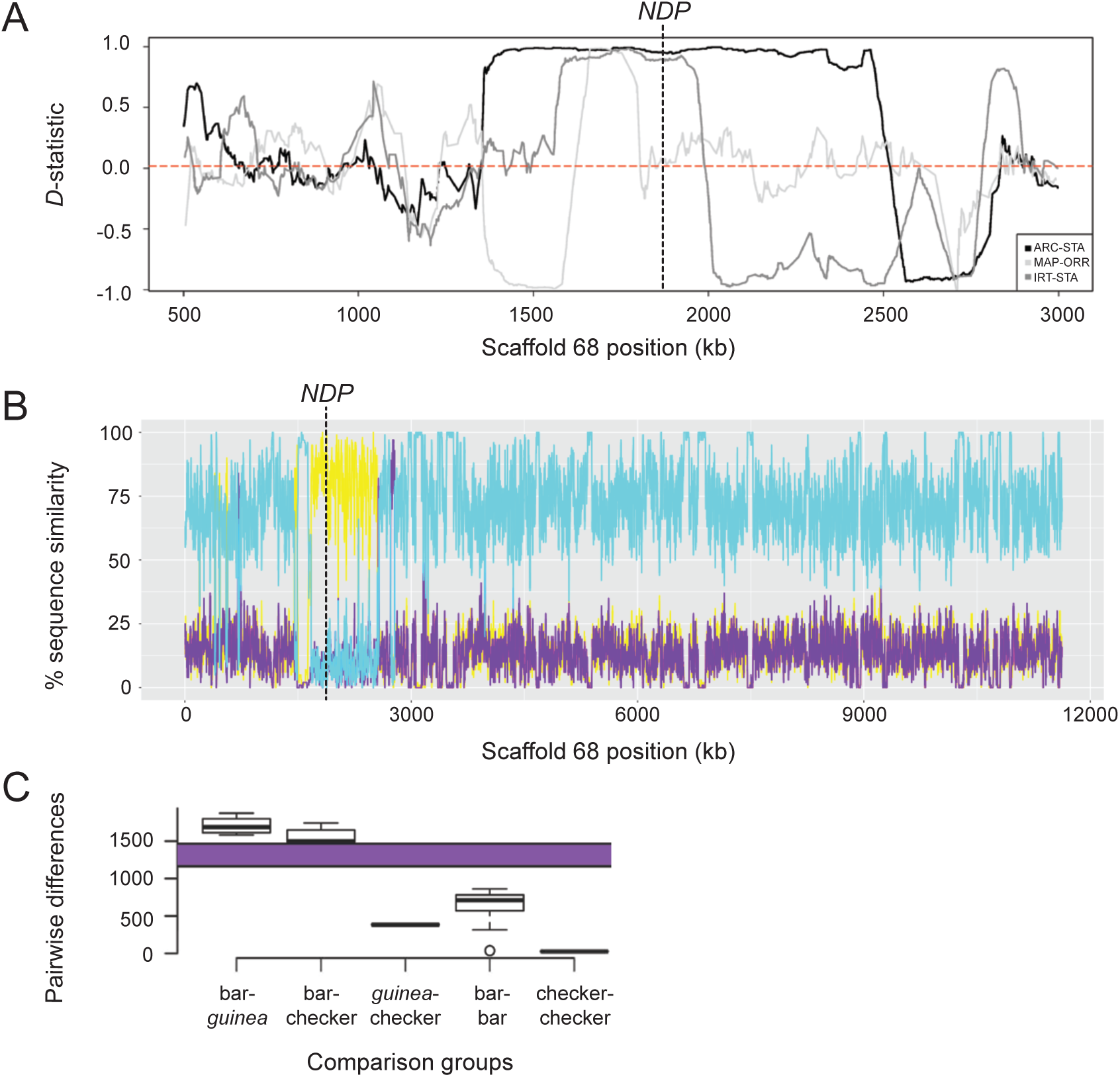
Signatures of introgression of the checker haplotype from *C. guinea* to *C. livia*. (A) ABBA-BABA test with *C. livia* (bar), *C. livia* (checker), *C. guinea*, and *C. palumbus* shows elevated D-statistic in the Scaffold 68 candidate region. Three representative ABBA-BABA tests are shown and dashed red line marks the genome-wide mean *D*-statistic for 10 X 10 different combinations of bar and checker birds (ARC-STA, MAP-ORR, IRT-STA are shown, see Methods). (B) HybridCheck shows sequence similarity between three pairwise comparisons: representative bar (Fer_VA), checker (ARC), and *C. guinea* individuals. (C) Expected (purple bar) and observed SNP differences in the minimal haplotype region for different pairwise comparisons between and among bar, checker, and *C. guinea*.

This unexpected similarity between *C. guinea* and checker *C. livia* in the pattern candidate region was further supported by sequence analysis using HybridCheck (Ward and van Oosterhout 2016). Outside of the candidate region, checker birds have a high sequence similarity to conspecific bar birds and low similarity to *C. guinea* (Fig. 4B). Within the candidate region, however, this relationship shows a striking reversal, and checker and *C. guinea* sequences are most similar to each other. In addition, although the genome-wide D-statistic was relatively low, the 95% confidence interval (CI) was greater than zero (0.021 to 0.022), providing further evidence for one or more introgression events from *C. guinea* into checker and T-check genomes. Unlike in many checker and T-check *C. livia*, we did not find additional copies of the CNV region in *C. guinea*. This could indicate that the CNV expanded in *C. livia*, or that the CNV is present in a subset of *C. guinea* but has not yet been sampled. Taken together, these patterns of sequence similarity and divergence support the hypothesis that the candidate checker haplotype in rock pigeons originated by introgression from *C. guinea*.

### Estimated divergence time among pattern locus haplotypes

While post-divergence introgression is an attractive hypothesis to explain the sequence similarity between checker *C. livia* and *C. guinea*, another formal possibility is that sequence similarity between these groups is due to incomplete lineage sorting. Therefore, to assess whether the minimal checker haplotype might have been present in the last common ancestor of *C. guinea* and *C. livia*, we measured single nucleotide differences among different alleles of the minimal haplotype and compared these counts to polymorphism rates expected to accumulate over the divergence time between *C. livia* and *C. guinea* (Fig. 4C, purple bar, see Methods).

We found that polymorphisms between bar *C. livia* and *C. guinea* approached the number expected to accumulate in 4-5 MY (1708±109 mean SNPs, Fig. 4C), but so did intraspecific comparisons between bar and checker *C. livia* (1564±99). In contrast, *C. guinea* and *C. livia* checker sequences had only 384±6 mean differences, significantly fewer than would be expected to accumulate between the two species (p < 2.2e-16, t-test). These results support an introgression event from *C. guinea* to *C. livia*, rather than a shared ancestral allele inherited from a common ancestor prior to divergence. Among 11 checker haplotype sequences we found only 26±8 mean differences.

We then estimated the age of the minimal checker haplotype following Voight et al. (2006). Using a recombination rate calculated for rock pigeon (Holt et al. 2017), the minimal checker haplotype is estimated to have been introgressed 857 (95% CI 534 to 1432) generations ago. Therefore, assuming 1-2 generations per year in *C. livia*, introgression events likely occurred well after the domestication of rock pigeons (~5000 years ago). The ranges of *C. livia* and *C. guinea* overlap in northern Africa (del Hoyo et al. 2017), so introgression events could have occurred in free-living populations. Alternatively, or perhaps additionally, multiple (but relatively similar) checker haplotypes could have been introgressed more recently by pigeon breeders. Once male hybrids are generated, this can be accomplished in just a few generations (Taibel 1949). This explanation is supported by lack of diversity among checker haplotypes, with only 26±8 mean differences, which is unusually low for the diversity typically observed in large, free-living pigeon populations (Shapiro et al. 2013). Additional *C. guinea* genome sequences will help to characterize allelic variation at this locus and resolve these possibilities.

### Introgression and pleiotropy

Adaptive traits can arise through new mutations or standing variation within a species, and a growing number of studies point to cross-species adaptive introgressions among vertebrates and other animals (Hedrick 2013; Harrison and Larson 2014; Zhang et al. 2016). In some cases, introgressed loci are associated with adaptive traits in the receiving species, including high-altitude tolerance in Tibetan human populations from Denisovans (Huerta-Sanchez et al. 2014), resistance to anticoagulant pesticides in the house mouse from the Algerian mouse (Song et al. 2011; Liu et al. 2015), and beak morphology among different species of Darwin’s finches (Lamichhaney et al. 2015). Among domesticated birds, introgressions are responsible for skin and plumage color traits in chickens and canaries, respectively (Eriksson et al. 2008; Lopes et al. 2016). Alleles under artificial selection in a domesticated species can be advantageous in the wild as well, as in the introgression of dark coat color from domestic dogs to wolves (Anderson et al. 2009) (however, color might actually be a visual marker for an advantageous physiological trait conferred by the same allele; Coulson et al. 2011).

In this study, we identified a putative introgression into *C. livia* from *C. guinea* that is advantageous both in artificial (selection by breeders) and free-living urban environments (sexual and natural selection). A change in plumage color pattern is an immediately obvious phenotypic consequence of the checker allele, yet other traits are linked to this pigmentation pattern. For example, checker and T-check pigeons have longer breeding seasons, up to year-round in some locations (Lofts et al. 1966; Murton et al. 1973), and *C. guinea* breeds year-round in most of its native range as well (del Hoyo et al. 2017). Perhaps not coincidentally, *NDP* is expressed in the gonad tissues of adult *C. livia* (MacManes et al. 2017) and the reproductive tract of other amniotes (Paxton et al. 2010). Abrogation of expression or function of *NDP* or its receptor *FZD4* is associated with infertility and gonad defects (Luhmann et al. 2005; Kaloglu et al. 2011; Ohlmann et al. 2012; Ohlmann and Tamm 2012). Furthermore, checker and T-check birds deposit less fat during normally reproductively quiescent winter months. In humans, expression levels of *FZD4* and the co-receptor *LRP5* in adipose tissue respond to varying levels of insulin (Karczewska-Kupczewska et al. 2016), and *LRP5* regulates the amount and location of adipose tissue deposition (Loh et al. 2015; Karczewska-Kupczewska et al. 2016). Therefore, based on its reproductive and metabolic roles in pigeons and other amniotes, *NDP* is a viable candidate not only for color pattern variation, but also for the suite of other traits observed in free-living (feral and wild) checker and T-check pigeons. Indeed, the potential pleiotropic effects of *NDP* raise the possibility that reproductive output and other physiological advantages are secondary or even primary targets of selection, and melanistic phenotypes are honest genetic signals of a cluster of adaptive traits controlled by a single locus.

Adaptive *cis*-regulatory change is emerging as an important theme in the evolution of vertebrates and other animals (Shapiro et al. 2004; Miller et al. 2007; Chan et al. 2010; Wittkopp and Kalay 2012; O’Brown et al. 2015). In some cases, the evolution of multiple regulatory elements of the same gene can fine-tune phenotypes, such as mouse coat color and trichome distribution in fruit flies (McGregor et al. 2007; Linnen et al. 2013). Cross-species introgression can result in the simultaneous transfer of multiple advantageous traits (Rieseberg 2011), and the potential role of *NDP* in both plumage and physiological variation in pigeons could represent a striking example of this process.

Wing pigmentation patterns that resemble checker are present in many wild species within and outside of Columbidae including *Patagioenas maculosa* (Spot-winged pigeon), *Spilopelia chinensis* (Spotted dove), *Geopelia cuneata* (Diamond dove), *Gyps rueppelli* (Rüppell’s vulture), and *Pygiptila stellaris* (Spot-winged antshrike). Based on our results in pigeons, *NDP* and its downstream targets can serve as initial candidate genes to ask whether similar molecular mechanisms generate convergent patterns in other species.

## MATERIALS & METHODS

### Ethics statement

Animal husbandry and experimental procedures were performed in accordance with protocols approved by the University of Utah Institutional Animal Care and Use Committee (protocols 10-05007, 13-04012, and 16-03010).

### DNA sample collection and extraction

Blood samples were collected in Utah at local pigeon shows, at the homes of local pigeon breeders, from pigeons in the Shapiro lab, and from ferals that had been captured in Salt Lake City, Utah. Photos of each bird were taken upon sample collection for our records and for phenotype verification. Tissue samples of *C. rupestris, C. guinea*, and *C. palumbus* were provided by the University of Washington Burke Museum, Louisiana State University Museum of Natural Science, and Tracy Aviary, respectively. Breeders outside of Utah were contacted by email or phone to obtain feather samples. Breeders were sent feather collection packets and instructions, and feather samples were sent back to the University of Utah along with detailed phenotypic information. Breeders were instructed to submit only samples that were unrelated by grandparent. DNA was then extracted from blood, tissue, and feathers as previously described (Stringham et al. 2012).

### Determination of color and pattern phenotype of adult birds

Feather and color phenotypes of birds were designated by their respective breeders. Birds that were raised in our facility at the University of Utah or collected from feral populations were assigned a phenotype using standard references (Levi 1986; Sell 2012).

### Genomic Analyses

BAM files from a panel of previously resequenced birds were combined with BAM files from 8 additional barless birds, 23 bar and 23 checker birds (22 feral, 24 domestics), a single *C. guinea*, and a single *C. palumbus*. SNVs and small indels were called using the Genome Analysis Toolkit (Unified Genotyper and LeftAlignAnd TrimVariants functions, default settings) (McKenna et al. (2010) *Genome Research*). Variants were filtered as described previously (Domyan et al. 2016) and the subsequent variant call format (VCF) file was used for pFst and ABBA-BABA analyses as part of the VCFLIB software library (https://github.com/vcflib) and VAAST (Yandell et al. 2011) as described previously (Shapiro et al. 2013).

pFst was first performed on whole-genomes of 32 bar and 27 checker birds. Some of the checker and bar birds were sequenced to very low coverage (~1X), so we were unable to confidently define the boundaries of the shared haplotype. To remedy this issue, we used the core of the haplotype to identify additional bar and checker birds from a set of birds that had already been sequenced to higher coverage (Shapiro et al. 2013). These additional birds were not included in the initial scan because their wing pattern phenotypes were concealed by other color and pattern traits that are epistatic to bar and check phenotypes. For example, the recessive red (*e*) and spread (*S*) loci produce a uniform pigment over the entire body, thereby obscuring any bars or checkers (Jones 1922; Hollander 1938; Sell 2012; Domyan et al. 2014). Although the major wing pattern is not visible in these birds, the presence or absence of the core checker haplotype allowed us to characterize them as either bar or checker/T-check. We then re-ran pFst using 17 bar and 24 checker/T-check birds with at least 8X mean read depth coverage and (Fig. 1B), and found a minimal shared checker haplotype of ~100 kb (Scaffold 68 position 1,702,691-1,805,600), as defined by haplotype breakpoints in a homozygous checker and a homozygous bar bird (NCBI BioSamples SAMN01057561 and SAMN01057543, respectively; BioProject PRJNA167554). pFst was also used to compare the genomes of 32 bar and 9 barless birds. New sequence data for *C. livia* are deposited in the NCBI SRA database under BioProject PRJNA428271 with the BioSample accession numbers SAMN08286792-SAMN08286844. (Submission of sequences for *C. guinea* and *C. palumbus* is in progress.)

### CNV breakpoint identification and read depth analysis

The approximate breakpoints of the CNV region were identified at Scaffold 68 positions 1,790,000 and 1,805,600 using WHAM in resequenced genomes of homozygous bar or checker birds with greater than 8x coverage (Kronenberg et al. 2015). For 12 bar, 7 checker, and 2 T-check resequenced genomes, Scaffold 68 gdepth files were generated using VCFtools (Danecek et al. 2011). Two subset regions were specified: the first contained the CNV and the second was outside of the CNV and was used for normalization (positions 1,500,000-2,000,000 and 800,000-1,400,000, respectively). Read depth in the CNV was normalized by dividing read depth by the average read depth from the second (non-CNV) region, then multiplying by two to normalize for diploidy.

### Taqman assay for copy number variation

Copy number variation was estimated using a custom Taqman Copy Number Assay (assay ID: cnvtaq1_CC1RVED; Applied Biosystems, Foster City, CA) for 94 birds phenotyped by wing pigment pattern category and 89 birds whose pigmentation was quantified by image analysis. After DNA extraction, samples were diluted to 5ng/μL. Samples were run in quadruplicate according to the manufacturer’s protocol.

### Quantification of pigment pattern phenotype

At the time of blood sample collection, the right wing shield was photographed (RAW format images from a Nikon D70 or Sony a6000 digital camera). In Photoshop (Adobe Systems Incorporated, San Jose, CA), the wing shield including the bar (on the secondary covert feathers) was isolated from the original RAW file. Images were adjusted to remove shadows and the contrast was set to 100%. The isolated adjusted wing shield image was then imported into ImageJ (imagej.nih.gov/) in JPEG format. Image depth was set to 8-bit and we then applied the threshold command. Threshold was further adjusted by hand to capture checkering and particles were analyzed using a minimum pixel size of 50. This procedure calculated the area of dark plumage pigmentation on the wing shield. Total shield area was calculated using the Huang threshold setting and analyzing the particles as before (minimum pixel size of 50). The dark area particles were divided by total wing area particles, and then multiplied by 100 to get the percent dark area on the wing shield. Measurements were done in triplicate for each bird, and the mean percentages of dark area for each bird were used to test for associations between copy number and phenotype using a non-linear least squares regression.

### qRT-PCR analysis of gene expression

Two secondary covert wing feathers each from the wing shields of 8 bar, 7 checker, and 8 T-check birds were plucked to stimulate feather regeneration for qRT-PCR experiments. Nine days after plucking, regenerating feather buds were removed, the proximal 5 mm was cut longitudinally, and specimens were stored in RNAlater (Qiagen, Valencia, CA) at 4°C for up to three days. Next, collar cells were removed, RNA was isolated, and mRNA was reverse-transcribed to cDNA as described previously (Domyan et al. 2014). Intron-spanning primers (see Table S1) were used to amplify each target using a CFX96 qPCR instrument and iTaq Universal Syber Green Supermix (Bio-Rad, Hercules, CA). Samples were run in duplicate and normalized to β-actin. The mean value was determined and results are presented as mean ± S.E. for each phenotype. Results were compared using a Wilcoxon Rank Sum test and expression differences were considered statistically-significant if p < 0.05.

### Allele-specific expression assay

SNPs in *NDP* and *EFHC2* were identified as being diagnostic of the bar or checker/T-check haplotypes from resequenced birds. Heterozygous birds were identified by Sanger sequencing in the minimal checker haplotype region (AV17 primers, see Table S1). Twelve checker and T-check heterozygous birds were then verified by additional Sanger reactions (AV54 for *NDP* and AV97 for *EFHC2*, see Table S1) to be heterozygous for the SNPs in *NDP* and *EFHC2*. PyroMark Custom assays (Qiagen) were designed for each SNP using the manufacturer’s software (Table S1). Pyrosequencing was performed on gDNA and cDNA derived from collar cells from 9-day regenerating feathers using a PyroMark Q24 instrument (Qiagen). Signal intensity ratios from the cDNA samples were normalized to the ratios from the corresponding gDNA samples to control for bias in allele amplification. Normalized ratios were analyzed by a Wilcoxon Rank Sum test and results were considered significant if p < 0.05.

### NDP genotyping and alignments

NDP exons were sequenced using primers in Table S1. Primers pairs were designed using the rock pigeon reference genome (Cliv_1.0) (Shapiro et al. 2013). PCR products were purified using a QIAquick PCR purification kit (Qiagen) and Sanger sequenced. Sequences from each exon were then edited for quality with Sequencher v.5.1 (GeneCodes, Ann Arbor, MI). Sequences were translated and aligned with SIXFRAME and CLUSTALW in SDSC Biology Workbench (http://workbench.sdsc.edu). Amino acid sequences outside of Columbidae were downloaded from Ensembl (www.ensembl.org).

### *D*-statistic calculations

Whole genome ABBA-BABA (https://github.com/vcflib) was performed on 10 X 10 combinations of bar and checker (Table S2) birds in the arrangement: bar, checker, *C. guinea*, *C. palumbus*. VCFLIB (https://github.com/vcflib) was used to smooth raw ABBA-BABA results in 1000-kb or 100-kb windows for whole-genome or Scaffold 68 analyses respectively. For each 10 X 10 combination. We calculated the average D statistic across the genome. These were then averaged to generate a whole genome average of D=0.0212, marked as the dotted line in Fig. 4A. Confidence intervals were generated via moving blocks bootstrap (Kunsch 1989). Block sizes are equal to the windows above, with D-statistic values resampled with replacement a number of times equal to the number of windows in a sample. In Figure 4A, three representative ABBA-BABA tests are shown for different combinations of bar and checker birds. The checker and bar birds used in each representative comparison are: ARC-STA, SRS346901 and SRS346887; MAP-ORR, SRS346893 and SRS346881; IRT-STA, SRS346892 and SRS346887 respectively.

### Haplotype phasing and HybridCheck analysis

VCF files containing Scaffold 68 genotypes for 16 bar, 11 homozygous checker, and 1 *C. guinea* were phased using Beagle version 3.3 (Browning and Browning 2007). VCFs were then converted to fasta format using vcf2fasta in vcf-lib (https://github.com/vcflib). HybridCheck (Ward and van Oosterhout 2016) (https://github.com/Ward9250/HybridCheck) was run to visualize pairwise sequence similarities between trios of bar, checker, and *C. guinea* sequences across Scaffold 68.

### Pairwise SNP comparisons

Phased VCF files for 16 bar, 11 homozygous checker, and 1 *C. guinea* were subsetted to the minimal checker haplotype region (positions 1,702,691-1,805,600) with tabix (Li 2011). The vcf-compare software module (VCFtools, Danecek et al. 2011) was used to run pairwise comparisons between bar, checker, and *C. guinea* birds (176 bar-checker, 16 bar-guinea, and 11 checker-guinea comparisons) as well as among bar and checker birds (120 bar-bar and 55 checker-checker comparisons). The total number of differences for each group was compared to the number of differences that are expected to accumulate during a 4-5 million year divergence time in a 102,909-bp region (the size of the minimal checker haplotype) with the mutation rate μ=1.42e-9 (Shapiro et al. 2013) using the coalescent equation: Time=#SNPs/(2xμx length of the region). The observed pairwise differences and the expected number of differences were evaluated with two-sample t-tests and all groups were considered statistically different from the 4-5 MY expectation (1169.05-1461.31). Standard deviations from the mean number of differences for each group were calculated in R: bar-*guinea*, 109; bar-checker, 99; bar-bar, 143; checker-*guinea*, 6; checker-checker, 8.

### Transcript amplification of barless allele of *NDP*

In order to determine whether the barless allele of *NDP* is transcribed and persists in the cell, or is degraded by the non-sense mediated decay (NMD) pathway, we designed a PCR assay to amplify *NDP* mRNA using intron-spanning primers (see Table S1). 4 barless, 2 bar, 2 checker, and 2 T-check birds were plucked to stimulate regeneration for NDP amplification. Feathers were harvested, RNA extracted, and cDNA synthesized as above. We detected expression of *NDP* in feather buds from barless feathers (n=4 feathers from a single individual). While not quantitative, expression was qualitatively similar to the levels of amplicons generated from other pattern phenotypes (n=2 for bar, checker, and T-check).

### *EFHC2* alignments

*EFHC2* exonic sequences from resequenced homozygous bar (n=16), homozygous check or T-check (n=11), barless (n=9), *Columba rupestris* (n=1), *Columba guinea* (n=1), and *Columba palumbus* (n=1) were extracted using the IGV browser (Thorvaldsdottir et al. 2013). Exon sequences for each group were translated using SIXFRAME in SDSC Biology Workbench (http://workbench.sdsc.edu). Peptide sequences were then aligned to EFHC2 amino acid sequences from other species downloaded from ensembl (http://www.ensembl.org) using CLUSTALW (Thompson et al. 1994) in SDSC Biology Workbench. Exon sequences from additional *C. livia* (n=17 checker or T-check and n=14 bar) and *C. guinea* (n=5) birds were determined by Sanger sequencing.

### Recombination rate estimation

Recombination frequency estimates were generated from a genetic map based an F2 cross of two divergent *C. livia* breeds, a Pomeranian Pouter and a Scandaroon (Domyan et al. 2016). Briefly, for genetic map construction, genotyping by sequencing (GBS) data were generated, trimmed, and filtered as described (Domyan et al. 2016), then mapped to the pigeon Cliv_1.0 pigeon genome assembly using Bowtie2 (Langmead and Salzberg 2012). Genotypes were called using Stacks (Catchen et al. 2011), and genetic map construction was performed using R/qtl (www.rqtl.org) (Broman et al. 2003). Pairwise recombination frequencies were calculated for all markers based on GBS genotypes. Within individual scaffolds, markers were filtered to remove loci showing segregation distortion (Chi-square, p < 0.01) or probable genotyping error. Specifically, markers were removed if dropping the marker led to an increased LOD score, or if removing a non-terminal marker led to a decrease in length of >10 cM that was not supported by physical distance. Individual genotypes with error LOD scores >5 (Lincoln and Lander 1992) were also removed. Pairwise recombination frequencies for markers retained in the final linkage map were used to estimate the age of the introgression event between *C. guinea* and *C. livia*.

### Minimal haplotype age estimation

The minimal haplotype age was estimated following Voight et al. (2006). We assume a star-shaped phylogeny, in which all samples with the minimal haplotype are identical to the nearest recombination event, and differ immediately beyond it. Choosing a variant in the center of the minimal haplotype, we calculated EHH, and estimated the age using the largest haplotype with a probability of homozygosity just below 0.25. Note that

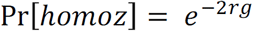
where r is the genetic map distance, and g is the number of generations since introgression / onset of selection. Therefore

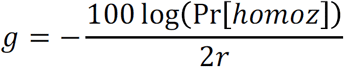

The confidence interval around g was estimated assuming

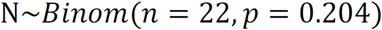

Here, N is a binomially distributed random variable for the number of samples that have not recombined to a map distance equal to 2r. Then, Pr[homoz] = N / 22. The probability that a sample has no recombination event within 2r of the focal SNP is p = (Pr[homoz | left] + Pr[homoz | right]) / 2 is derived from the data. Both left and right of the focal SNP we chose the end of the haplotype at the first SNP which brought Pr[homoz] < 0.25.

## ACKNOWLEDGEMENTS

We thank past and present members of the Shapiro lab for assistance with sample collection and processing; members of the Utah Pigeon Club and National Pigeon Association for sample contributions; and Gene Hochlan, Gary Young, and Robert Mangile for critical discussions and advice. Mr. Hochlan also generously provided feather samples from *C. guinea* that helped us assess feasibility of the introgression study. We thank Fred Adler, Brett Boyd, Elena Boer, Robert Greenhalgh, J.J. Horns, Christopher Leonard, Raquel Maynez, Jon Seger, and Scott Villa for technical assistance and advice. Dale Clayton and Sarah Bush generously provided field-collected tissue samples of *C. guinea* and *C. palumbus* for whole-genome sequencing. We thank Safari West (Santa Rosa, CA), the Louisiana State University Museum of Natural Science, nad the University of Washington Burke Museum for additional *C. guinea* tissue samples (museum accessions 95045 JK 00 179, 101559 BCA 523, and 119004 EEM 979). This work was supported by the National Science Foundation (CAREER DEB-1149160 to M.D.S.; GRF 1256065 to A.I.V. and R.B.; and DEB-1342604 to K.P.J.) and the National Institutes of Health (R01GM115996 to M.D.S., R01GM104390 to M.Y.; fellowships T32GM007464 to Z.K. and R25CA057730 to R.J.B.). E.T.M. is a fellow of the Jane Coffin Childs Memorial Fund for Medical Research. The funders had no role in study design, data collection and analysis, decision to publish, or preparation of the manuscript. We acknowledge a computer time allocation from the Center for High Performance Computing at the University of Utah.

## Author contributions

A.I.V., Z.K., C.D.H., M.Y., and M.D.S. designed research; A.I.V., Z.K., R.B., and E.M. performed research; A.I.V., Z.K., R.B., E.M., R.J.M., E.T.M., E.J.O., and M.D.S. analyzed data; K.P.J. contributed biological samples and genome sequence data; and A.I.V. and M.D.S. wrote the paper with input from the other authors.

---

**Fig. S1.**
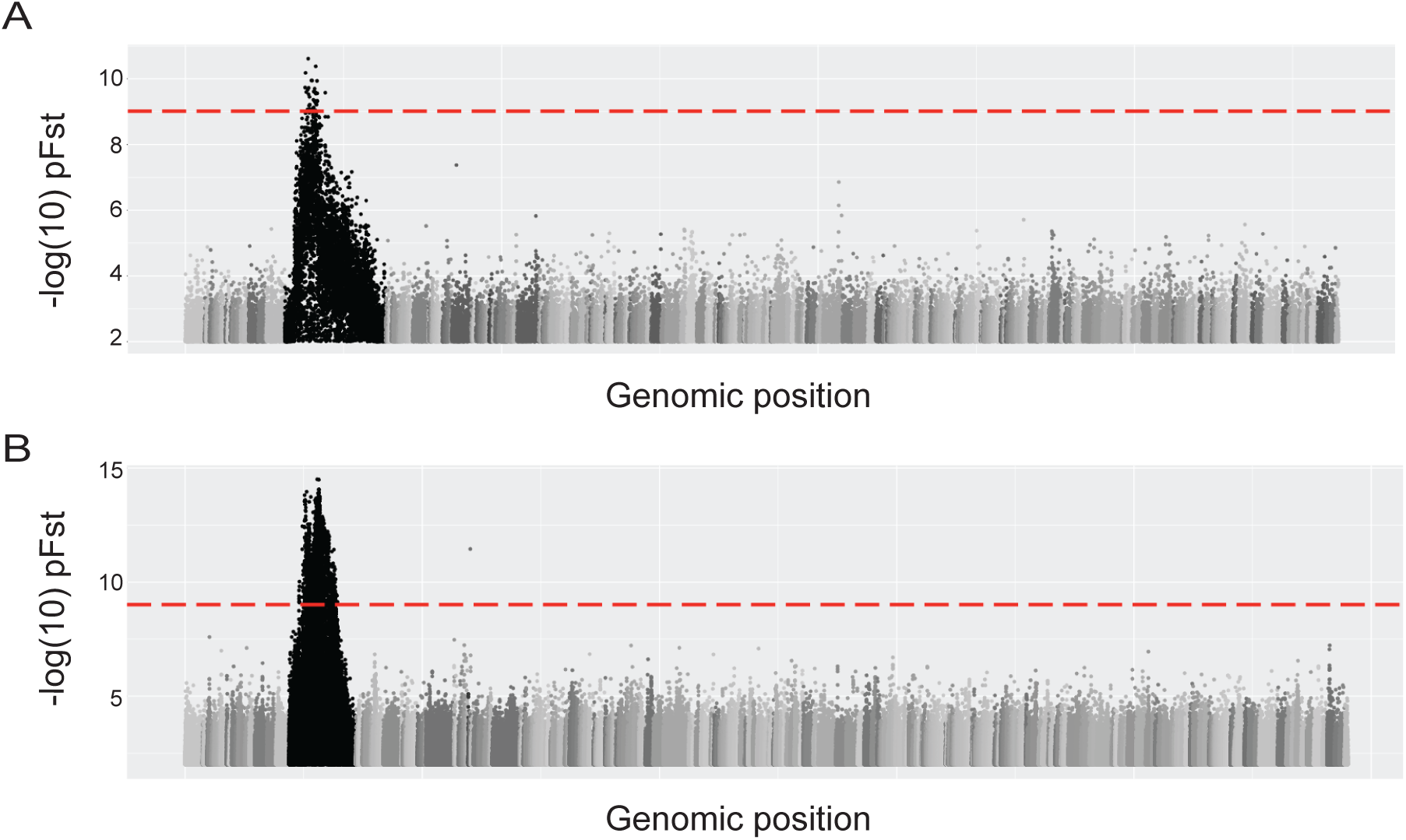
Whole genome pFst comparisons to identify a candidate genomic region differentiated between birds with different wing pattern phenotypes. (A) Whole genome pFst comparing 32 bar and 27 checker and T-check birds. (B) Whole genome pFst comparing 32 bar and 9 barless birds.

**Fig. S2.**
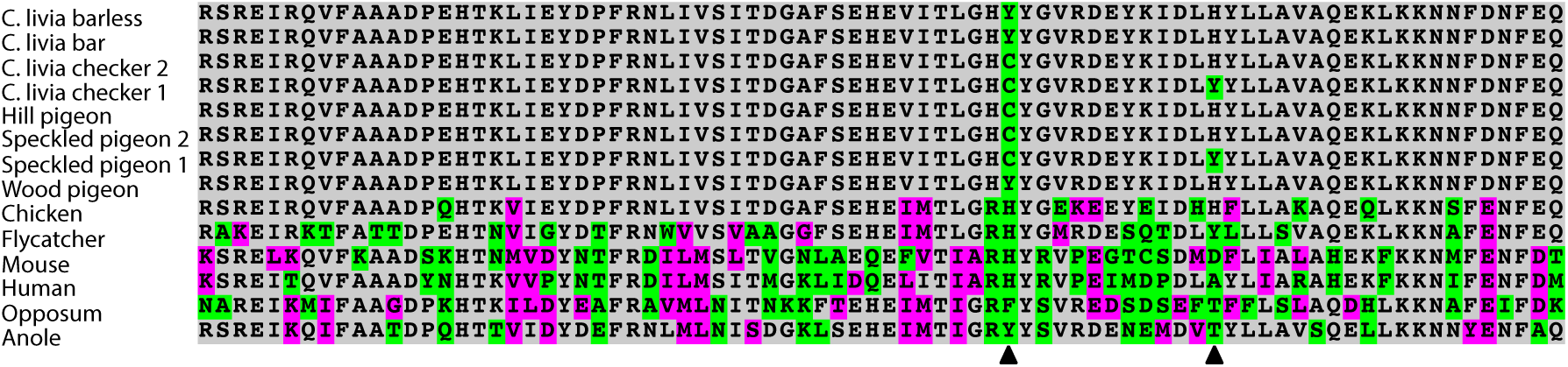
EFHC2 amino acid sequences of pigeons and other amniotes (residues 525–604). Variable amino acid residues are marked in magenta (similar residues) and green (different residues). Checker *C. livia, C. rupestris*, and *C. guinea* share 572C while bar *C. livia* are fixed for 572Y (left arrowhead). Checker *C. livia* and *C. guinea* are polymorphic for 584H/Y (right arrowhead).

**Fig. S3.**
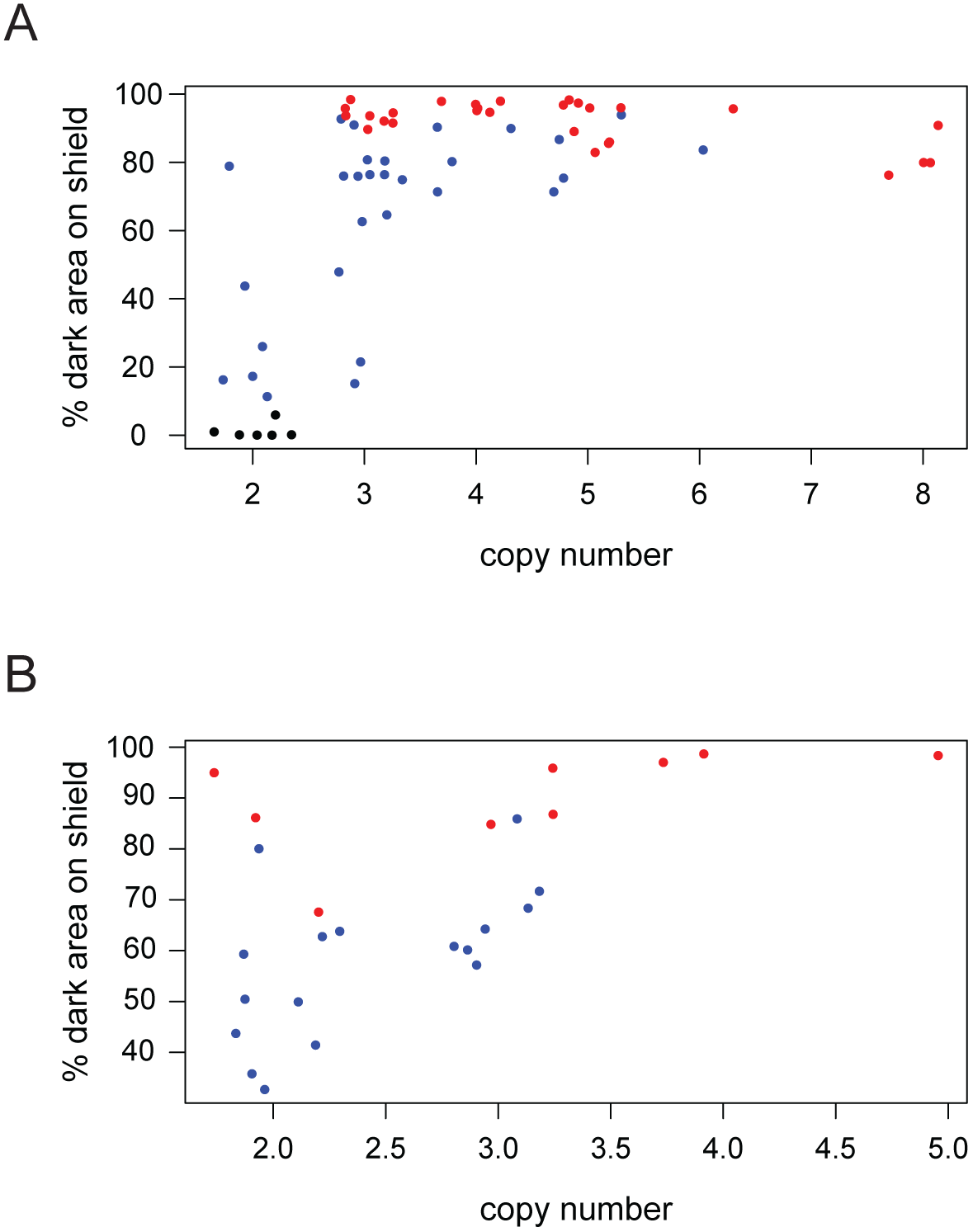
CNV is associated with darker wing shield pigmentation. CNV and phenotype quantification for (A) domestic breeds (n=58) and (B) wild-caught ferals (n=26), parsed from data in Fig. 2C.

**Fig. S4.**
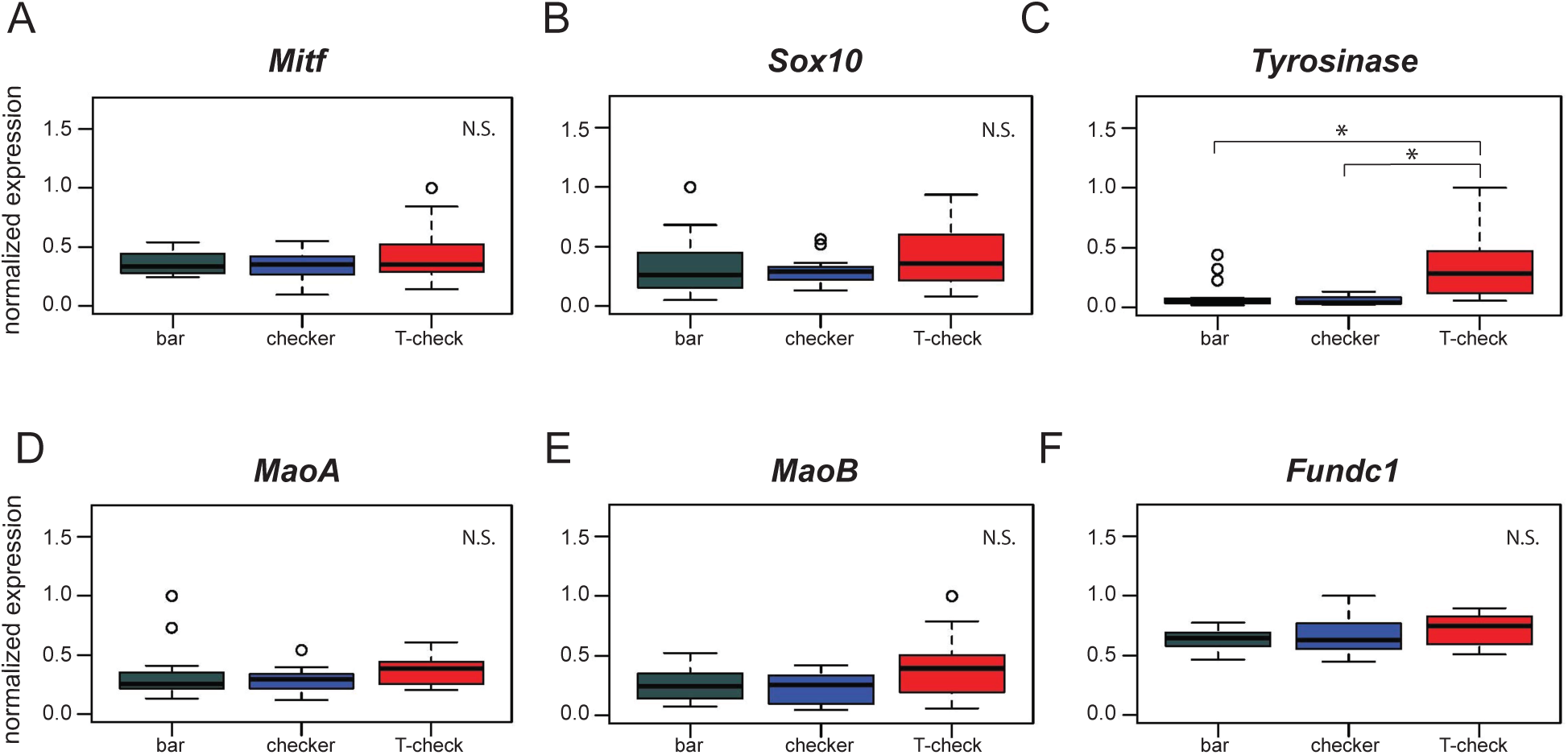
Expression of genes involved in pigmentation and genes in the candidate region. Expression levels of *Mitf* (A), *Sox10* (B), *MaoA* (D), *MaoB* (E), and *Fundc1* (F) are indistinguishable across phenotypes. (C) *Tyrosinase* shows increased expression in T-check birds relative to bar (p=2.4e-04) and checker birds (p=3.8e-05). Boxes span the first to third quartiles, bars extend to minimum and maximum observed values, black line indicates median. Expression values are analyzed by Pairwise Wilcoxon test (p-value adjustment method: fdr).

**Fig. S5.**
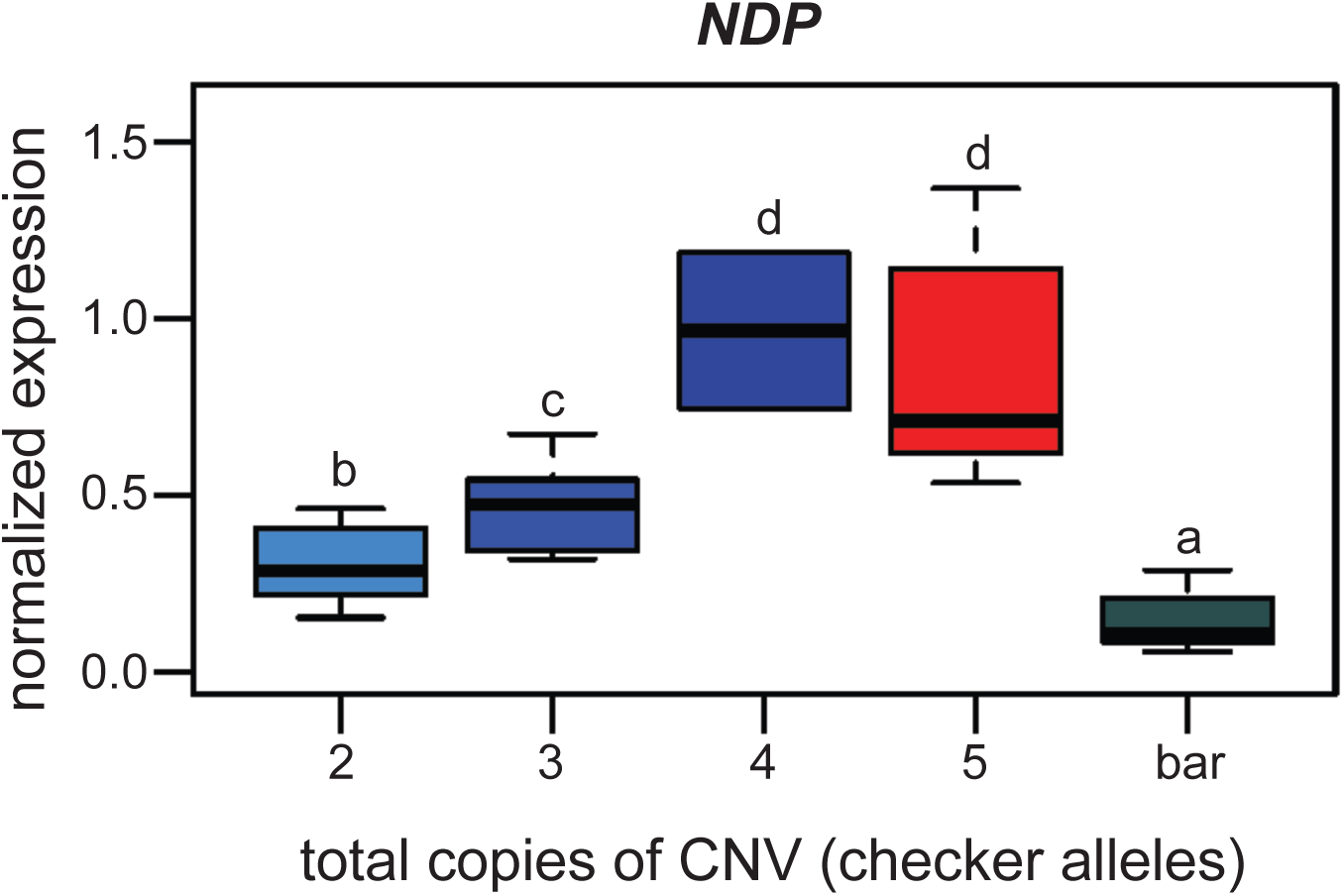
*NDP* expression varies by copy number and phenotype. qRT-PCR expression assay for *NDP* (Fig. 3A) is parsed by copy number in the CNV region. All checker (blue) and T-check (red) birds, except for the single individual with four total copies (dark blue), are heterozygous for bar. Increase in *NDP* expression is correlated with increasing numbers of copies of the CNV region. Boxes span the first to third quartiles, bars extend to minimum and maximum observed values, black line indicates median. Different letters indicate significant pairwise differences. Pairwise Wilcoxon test (p-value adjustment method: fdr) results by copy number: 2–3 copies, p=0.03788; 2–4 copies, p=0.04938; 2–5 copies, p=0.00015; 2 copies-bar, p=0.00432; 3–4 copies, p=0.03788; 3–5 copies, p=0.00122; 3 copies-bar, p=1.9e-06; 4–5 copies, p=0.48485; 4 copies-bar, p=0.02179; 5 copies-bar, p=1.9e-06.

**Fig. S6.**
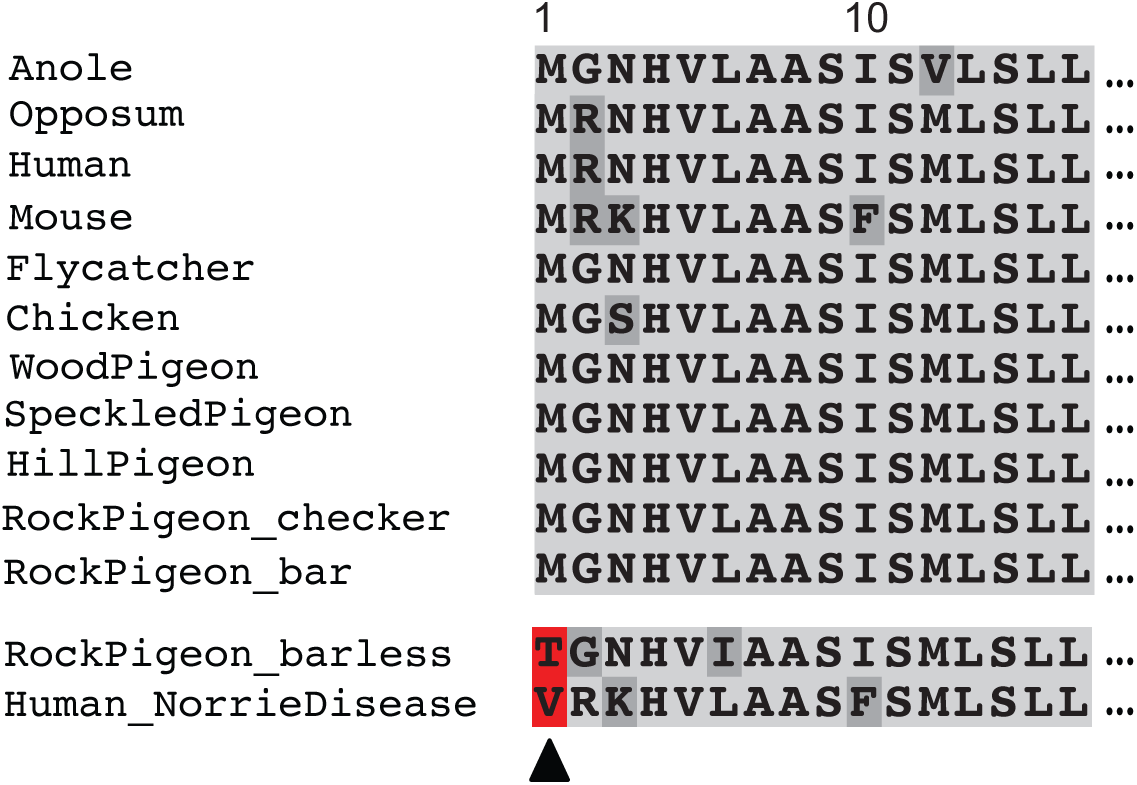
Barless pigeons have a nonsense mutation at the highly-conserved translation start site of *NDP*. Barless rock pigeons are homozygous for a nonsense mutation that truncates the amino terminus of *NDP* to 13 amino acids; the same amino acid position is affected by a mutation in two human families with hereditary blindness (red, bottom of alignments).

**Fig. S7.**
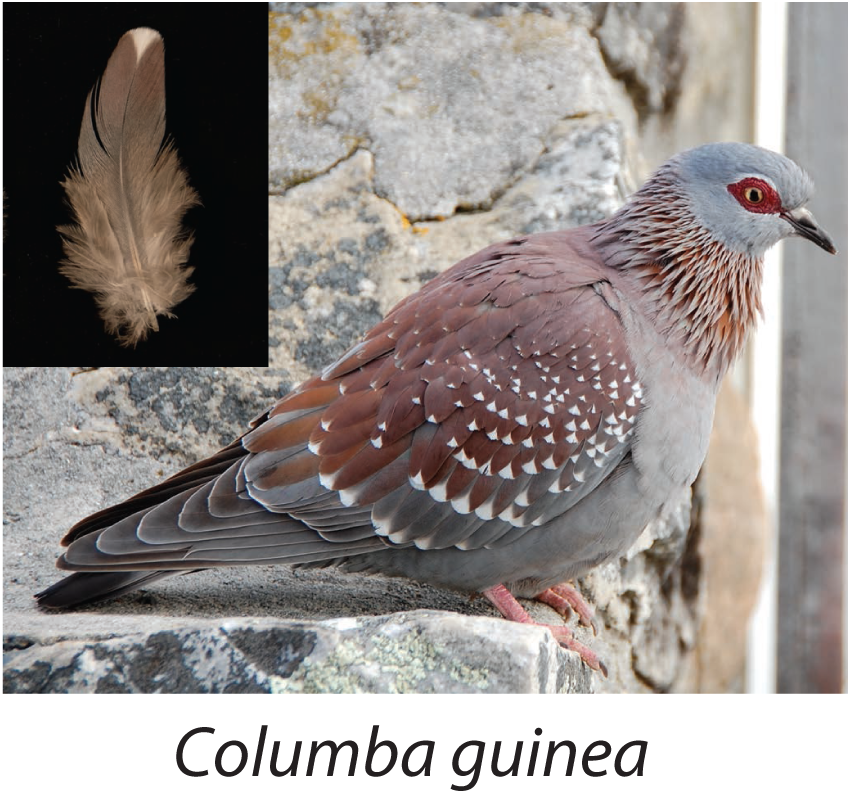
Speckled pigeon (*Columba guinea*). Photo courtesy of Kjeuring (CC BY 3.0 license, https://creativecommons.org/licenses/by/3.0/legalcode). Photo cropped from “speckled pigeon *Columba guinea* Table Mountain Cape Town,” https://en.wikipedia.org/wiki/Speckled_pigeon#/media/File:Speckledpigeon.JPG. Inset feather image by the authors.

